## Supplementary Information for "Generating information-dense promoter sequences with optimal string packing"

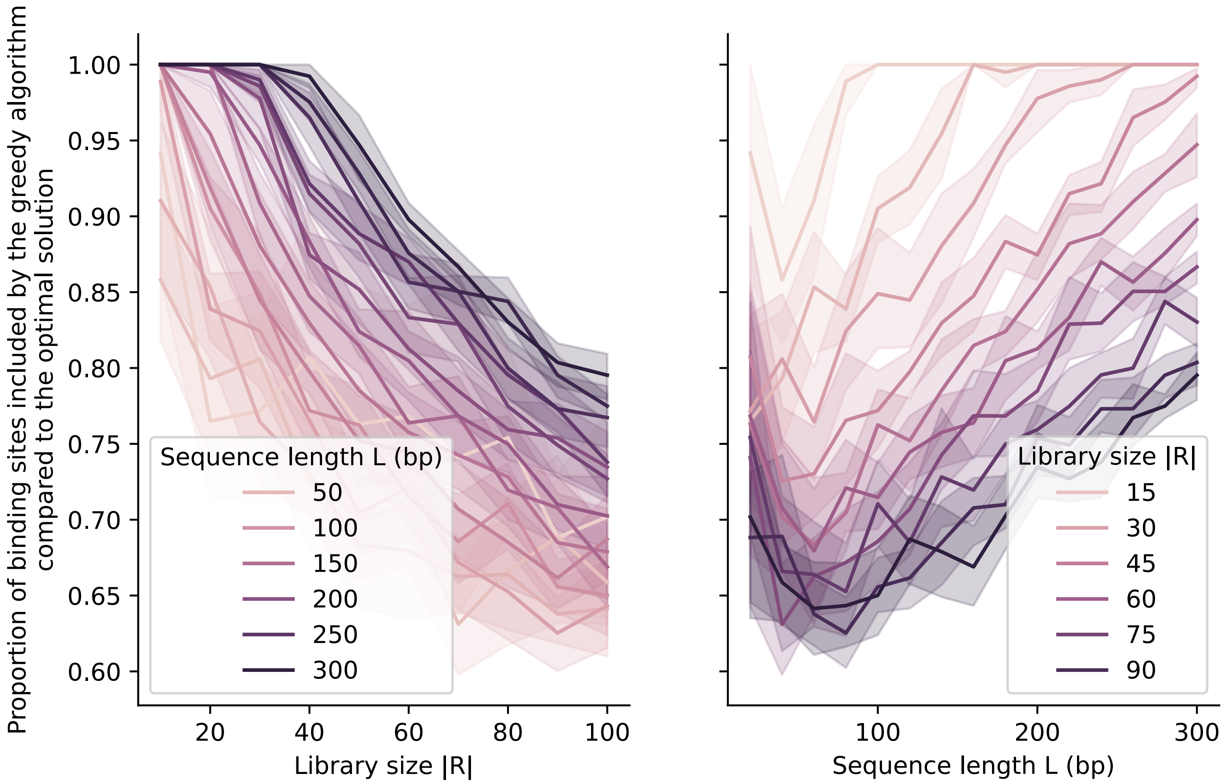


**Figure S1.** Performance of the greedy approximation algorithm. We plotted the number of binding sites that the approximation algorithm managed to fit into a sequence of a given length, *L*, normalized by the optimal solution. We repeated each scenario 10 times with random binding sites of uniform random lengths between 5 and 15 base pairs. Shaded regions represent the bootstrapped 95% confidence interval around the mean.

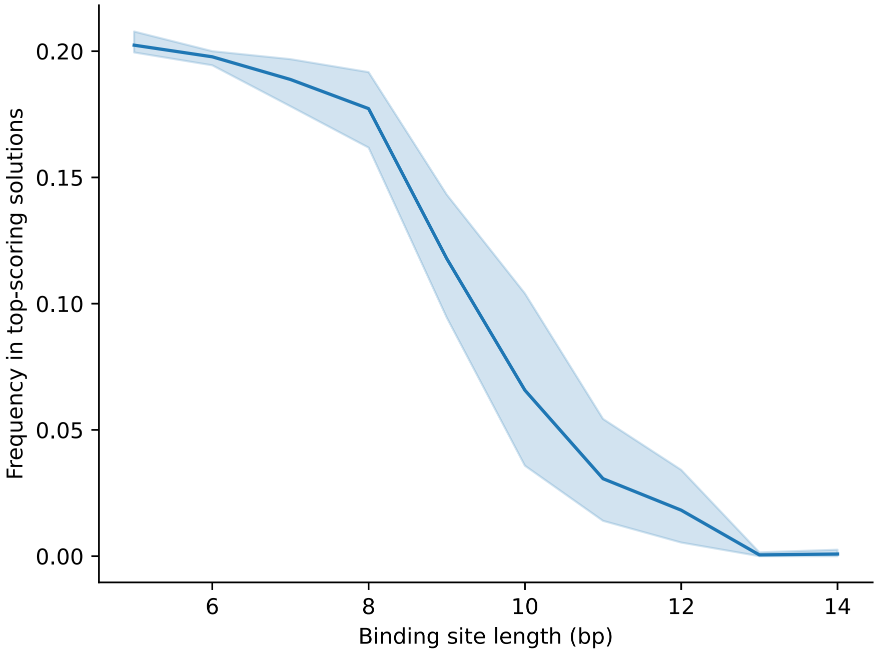


**Figure S2.** Frequency of binding site usage as a function of binding site length. Ten libraries were randomly generated with 10 binding sites each, where there is one binding site of each length from 5 to 14 base pairs. All of the top-scoring solutions for a sequence length $L = 50$ were generated every time: for one of the 10 libraries, 4 binding sites were able to be fit at best, with 7184 ways to do so. For the nine others, 5 binding sites were able to be fit at best, with 26, 72, 150, 184, 206, 278, 418, 664, 762 ways to do so. The shaded region represents the bootstrapped 95% confidence interval around the mean.

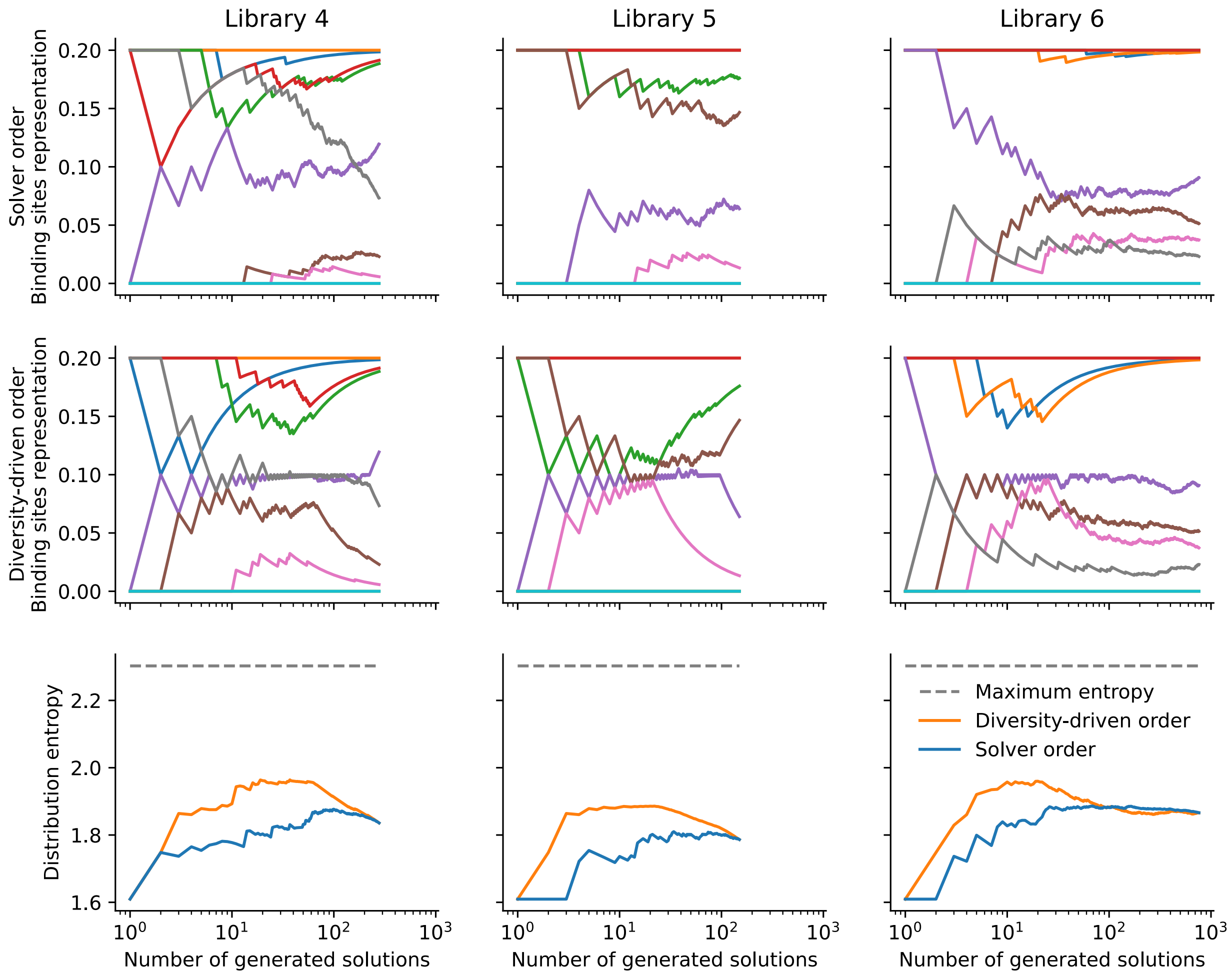


**Figure S3.** The distribution control strategy (i.e., diversity-driven order) applied to heavily biased libraries where binding sites have different sizes. Every library is made of 10 randomly generated binding sites, one of size 5 base pairs, one of size 6, etc., until size 14. As the full distribution attests (rightmost point of the graphs), some binding sites are present in almost all top-scoring solutions, while some others are present in none. Despite this, the diversity-driven order approach generally produces higher entropy distributions.
